# Reconstruction of the binding pathway of an anti-HIV drug, Indinavir, in complex with the HTLV-1 protease by Unaggregated Unbiased Molecular Dynamics simulation

**DOI:** 10.1101/2021.09.09.459615

**Authors:** Farzin Sohraby, Hassan Aryapour

**Affiliations:** Department of Biology, Faculty of Science, Golestan University, Gorgan, Iran

**Author notes:** Corresponding author. Department of Biology, Faculty of Science, Golestan University, Gorgan, Iran.

**Keywords:** Human T-cell Leukemia Virus 1 (HTLV-1) virus, Indinavir, HTLV-1 protease, Unaggregated Unbiased Molecular Dynamics (UUMD), Binding pathway

## Abstract

Retroviruses are a growing concern for the health of human beings, and one of the dangerous members of this family is the Human T-cell Leukemia Virus 1 (HTLV-1) virus. It has affected more than 20 million people so far, and since there are no registered treatments against it yet, urgent treatment solutions are needed. One of the most promising drug targets to fight this virus is the protease enzyme of the virus’s protein machinery. In this study, by utilizing a computational method called Unaggregated Unbiased Molecular Dynamics (UUMD), we reconstructed the binding pathway of a HTLV-1 protease inhibitor, Indinavir, to find the details of the binding pathway, the influential residues, and also the stable states of the binding pathway. We achieved the native conformation of the inhibitor in 6 rounds, 360 replicas by performing over 4 micro-seconds of UMD simulations. We found 3 Intermediate states between the solvated state and the native conformation state in the binding pathway. We also discovered that aromatic residues such as Trp98 and Trp98′, catalytic residues Asp32 and Asp32′, and the flap region’s residues have the most influential roles in the binding pathway and also have the most contribution to the total interaction energies. We believe that the details found in this study would be a great guide for developing new treatment solutions against the HTLV-1 virus by inhibiting the HTLV-1 protease.

## Introduction

Retroviruses are among the most dangerous viruses that have always been a threat to human beings’ lives. The RNA-enveloped Human T-cell Leukemia Virus 1 (HTLV-1) virus, which is associated with adult T-cell leukemia (ATL) and an inflammatory disease syndrome called HTLV-1-associated myelopathy/tropical spastic paraparesis (HAM/TSP) [1-3], is one of the most dangerous retroviruses known to man [4-6]. It has infected more than 20 million people worldwide, and until this time, there is no registered treatment against this virus [3].

Unlike the HTLV-1 virus, there are many drugs to treat HIV that can inhibit most of the vital protein targets of the virus’s protein machinery, such as the HIV protease [7]. Repurposing or optimizing existing anti-HIV treatments to fight the HTLV-1 virus can be an excellent strategy for drug design since the two viruses have many similarities. In 2015, Kuhnert et al. tested existing HIV protease inhibitors on HTLV-1 protease to repurpose the existing drugs to treat the HTLV-1 virus [8]. However, among all of the HIV protease inhibitors, only Indinavir showed promising results compared to the others, with an inhibitory constant of 3.5 µM. Indinavir is one of the most effective HIV protease inhibitors, which has led to a significant advancement in the treatment of HIV infection [9]. It is a potent inhibitor of HTLV-1 protease, and optimizing and refining it towards better inhibition activity can be a good strategy to design an efficacious drug against the HTLV-1 virus. Optimizing this inhibitor partly requires a complete understanding of the structural details of the binding pathway of the inhibitor in complex with its target protein.

Structure-based drug discovery (SBDD) by computational tools is a rapidly growing area with promising capabilities which can be used for situations like this that urgent treatment solutions are needed [10-12]. Fast and accurate methods such as Unbiased Molecular Dynamics (UMD) simulations can also be utilized to unravel the binding pathways of small molecule inhibitors to evaluate the efficacy of potential inhibitors and understand their inhibitory mechanisms against a protein target [13-21]. The role of different factors in the binding mechanism has to be thoroughly analyzed and understood, and in the past few years, we have developed an efficient method called Unaggregated Unbiased Molecular Dynamics (UUMD) simulation for this purpose [15]. By introducing repulsive forces between the virtual interaction sites (VIS), which are located on the ligand molecules, we can insert a high concentration of ligand molecules into the simulation box. This approach increases the chance of molecules to sample the target protein’s surface and find the binding site with the correct orientation and reach the native binding pose.

In this study, by utilizing the UUMD method, we tried to reconstruct the binding pathway of Indinavir to the HTLV-1 protease and understand the details and the factors involved in the binding pathway. This study can benefit researchers in SBDD to better understand the details of the inhibition mechanisms of this inhibitor, which will then facilitate the design of more selective and more effective inhibitors to find an immediate treatment for the HTLV-1 associated diseases.

## Methods

For this study, the crystallography structure of the HTLV-1 protease (PDB ID: 3WSJ) was obtained from the PDB database [22]. Firstly, the structure was cleaned, and only the protein and the inhibitor were kept, and other molecules such as water molecules were deleted. Then, the apo form of the protein was put in the center of the simulation box with a minimum distance of 1.5 nm to the edges. Then, 16 molecules of Indinavir were added to the box randomly. After that, the system was solvated with the TIP3P water model [23], and sodium and chloride ions were added to produce a neutral physiological salt concentration of 150 mM. Each system was energy minimized using the steepest descent algorithm until the Fmax was less than 10 kJ.mol^-1^.nm^-1^. All of the covalent bonds were constrained using the Linear Constraint Solver (LINCS) algorithm [24] to maintain constant bond lengths. The long-range electrostatic interactions were treated using the Particle Mesh Ewald (PME) method [25], and the cut-off radii for Coulomb and Van der Waals short-range interactions were set to 0.9 nm. The modified Berendsen (V-rescale) thermostat [26] and the Parrinello–Rahman barostat [27] were applied for 100 ps of NVT and 300 ps of NPT equilibration runs, respectively, to keep the system in stable environmental conditions (310 K, 1 Bar). All of the MD simulations were done by GROMACS 2018 package [28] and OPLS-AA force field [29]. Finally, the simulations were carried out under the periodic boundary conditions (PBC), set at XYZ coordinates to ensure that the atoms had stayed inside the simulation box, and the subsequent analyses were then performed using GROMACS utilities, VMD [30] and USCF Chimera, and also the plots were created using Daniel’s XL Toolbox (v 7.3.2) add-in[31]. The free energy landscapes were rendered using Matplotlib[32]. In addition, to estimate the binding free energy, we used the g_mmpbsa package[33]. The partial charges and force field parameters of the inhibitor molecule were assigned by ACEPYPE [34] with the default setting. The pattern and the details of the VISs were set the same as the previous studies [15], the values of the σ and ε parameters for VISs were set to 0.83 nm and 0.1 kJ.mol^-1^, respectively.

## Results & Discussion

As explained in a previous study [15], the route for reconstructing the binding pathway of a ligand to the target protein is divided into separate stages. In each stage, the simulations are carried out by using short replicas. For example, instead of performing one long simulation for the sampling stage, 50 replicas with a duration time of 10 ns each were performed. Every 10 ns replica consists of 1000 frames, saving frames every 10 ps. After finding a successful replica, the best frame from the simulation was chosen and extracted to be the starting point of the next stage. The flowchart of the stages is shown in Figure 1. The criteria for choosing the best frame for the first step may be different from the other steps. In this study, in the first stage, the criteria for choosing the best frame were the closeness of a molecule to the binding pocket and also the orientation of that molecule at the entrance of the binding pocket, whereas, in the other stages, the criterion was reaching lower Root Mean Square Distance (RMSD). The results and the details of each stage and the criteria will be explained in the following sections.

**Figure 1.**
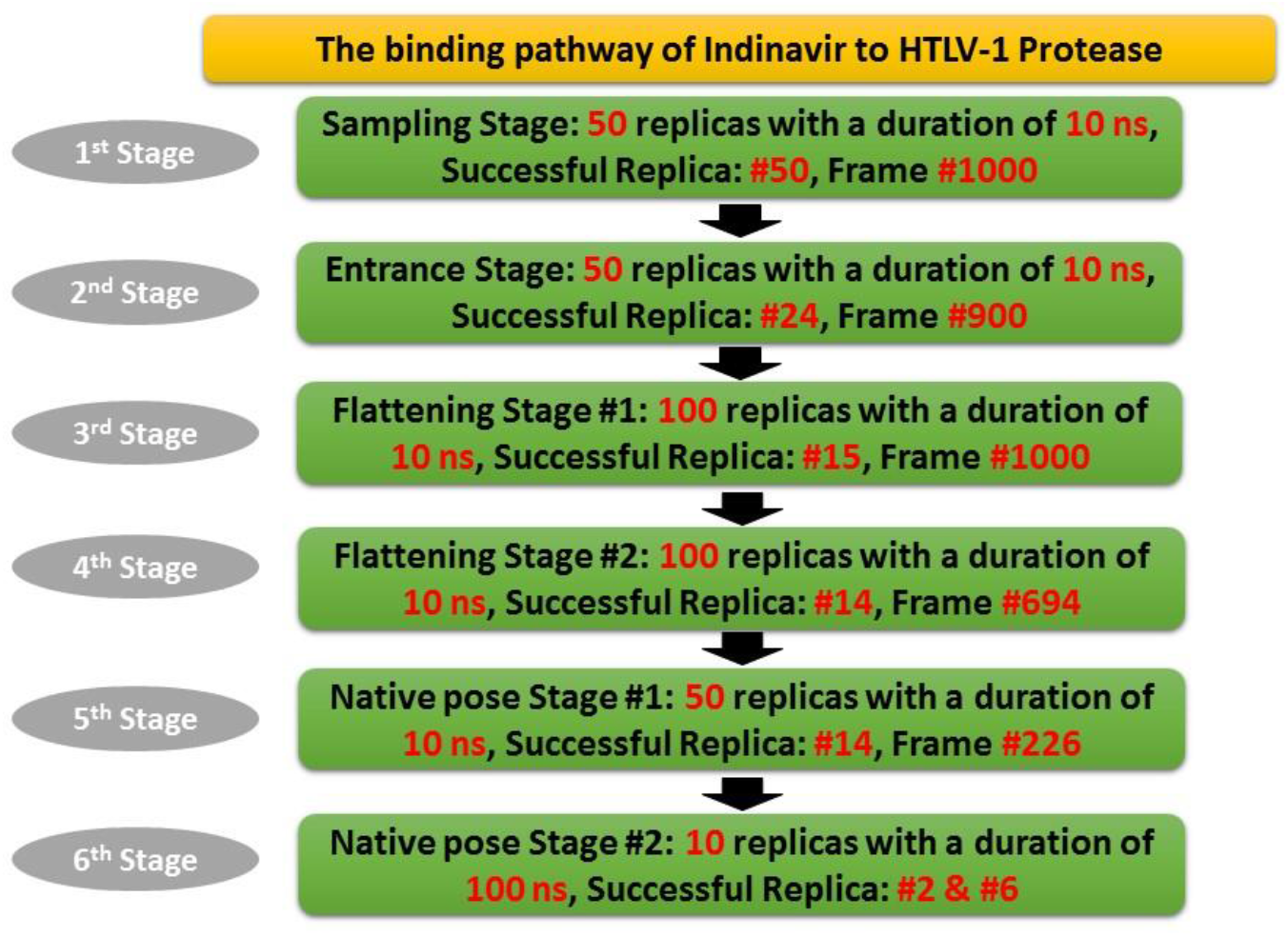
The flowchart of the reconstruction of Indinavir’s binding pathway to HTLV-1 protease.

In the first stage, the sampling stage, 16 Indinavir molecules were present in the simulation box with an approximate concentration of 45 mM. The simulation box with the Indinavir molecules inserted is shown in Fig. 2a. There was also a repulsive force between the Indinavir molecules that stops them from aggregating to each other. This repulsive force was applied through the VISs, which were located on the bonds between heavy atoms of the Indinavir molecules)Fig. 2e(. In this stage, 16 molecules were solvated in water and free to roam around the simulation box and sample the protein’s surface to find the binding pocket. During the 10 ns of MD simulation in each replica, some molecules got attracted to the protein’s surface, but others remained solvated. Each replica was thoroughly analyzed and inspected to find the best replica and the best frame. The criteria for choosing the best frame were the closeness of one of the 16 molecules to the binding pocket with appropriate orientation. Eventually, in replica #50, appropriate orientation of an Indinavir molecule was achieved, chosen, and selected as the best frame and used to perform the next stage’s replicas (Fig. 2b and 2c). The high concentration of ligand molecules and the repulsive force are only applied in the first stage. Finding the binding pocket by one of the ligand molecules marks the end of the first stage.

**Figure 2.**
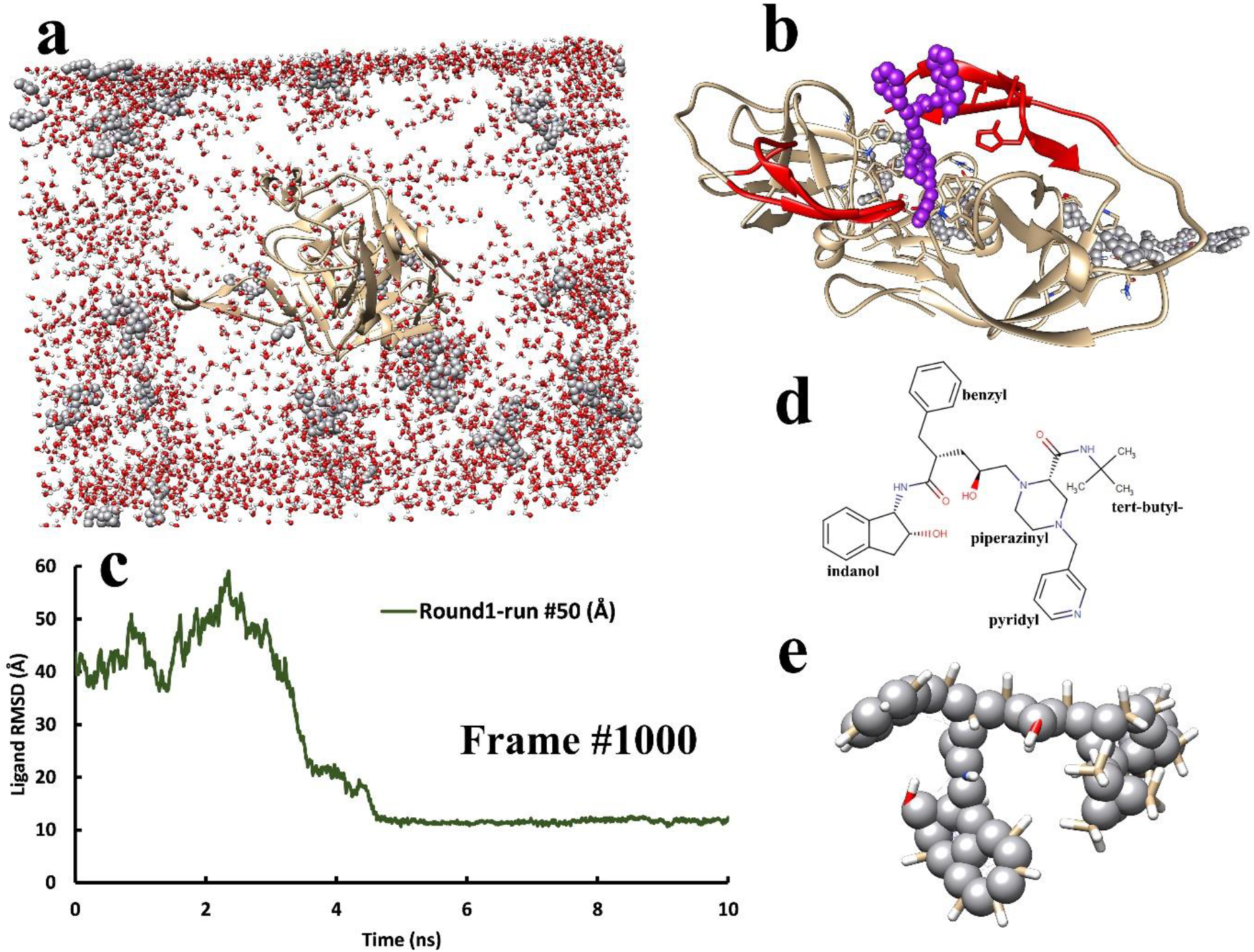
The first stage of the UUMD simulations, The sampling stage. **a**, The simulation system where the HTLV-1 protease and 16 Indinavir molecules were inserted randomly into the simulation box, which was then solvated by water molecules. **b**, The best frame of replica #50 in the 1^st^ stage in which one of the 16 Indinavir molecules interacted with the flap region (red) at the entrance of the binding site. This frame was chosen as the starting point of the next stage. c, The RMSD values of the chosen ligand molecule during the 10 ns production run, replica #50. d, The chemical structure of Indinavir and its various chemical groups. **e**, The position of the VISs on the structure of Indinavir were located on the bonds of heavy atoms of the Indinavir molecule.

The binding pathway of Indinavir to the HTLV-1 protease is not a straight path. The molecule interacts with different protein regions in each stage using appropriate chemical groups (Fig. 2d). Moreover, the orientation of the molecule was also essential. The orientation of the co-crystallized Indinavir molecule was considered as the reference to choose the best frame. In the #50 replica in the first stage, a molecule started interacting to the flap region with the correct orientation, shown in purple (Fig. 2b).

For the 2^nd^ stage, the entrance stage, we also performed 50 independent replicas lasting 10 ns each. From this stage onwards, only one Indinavir molecule was present in the simulation system, selected based on its proper orientation (Fig.2d). The other molecules and also the VISs with the repulsive force were omitted from the MD simulations. Out of 50 replicas, replica #24 was the most successful run, and the best frame was chosen from this replica, Frame #900. In the entrance stage, the Indinavir molecule had to interact more with the inner parts of the binding site and interact less with the flap region. The flap region covers the top parts of the binding pocket and acts as a gate for the entrance of the Indinavir molecule. In this stage, it was observed that Indinavir could engage with the flap region and stay connected. Although in the successful replica, replica #24, it was observed that the Indinavir molecule started to interact with the aromatic residues such as Trp98 and Trp98′ from both chains, the benzyl group of Indinavir was still interacting with the flap region’s residues (Fig. 3b). It was able to make π-stacking interactions with its indanol and pyridyl groups. The interaction with the binding pocket’s inner residues was considered the entrance of the molecule to the binding site. The frame shown in Fig. 3b was used as the starting frame of the MD simulations of the next stage.

**Figure 3.**
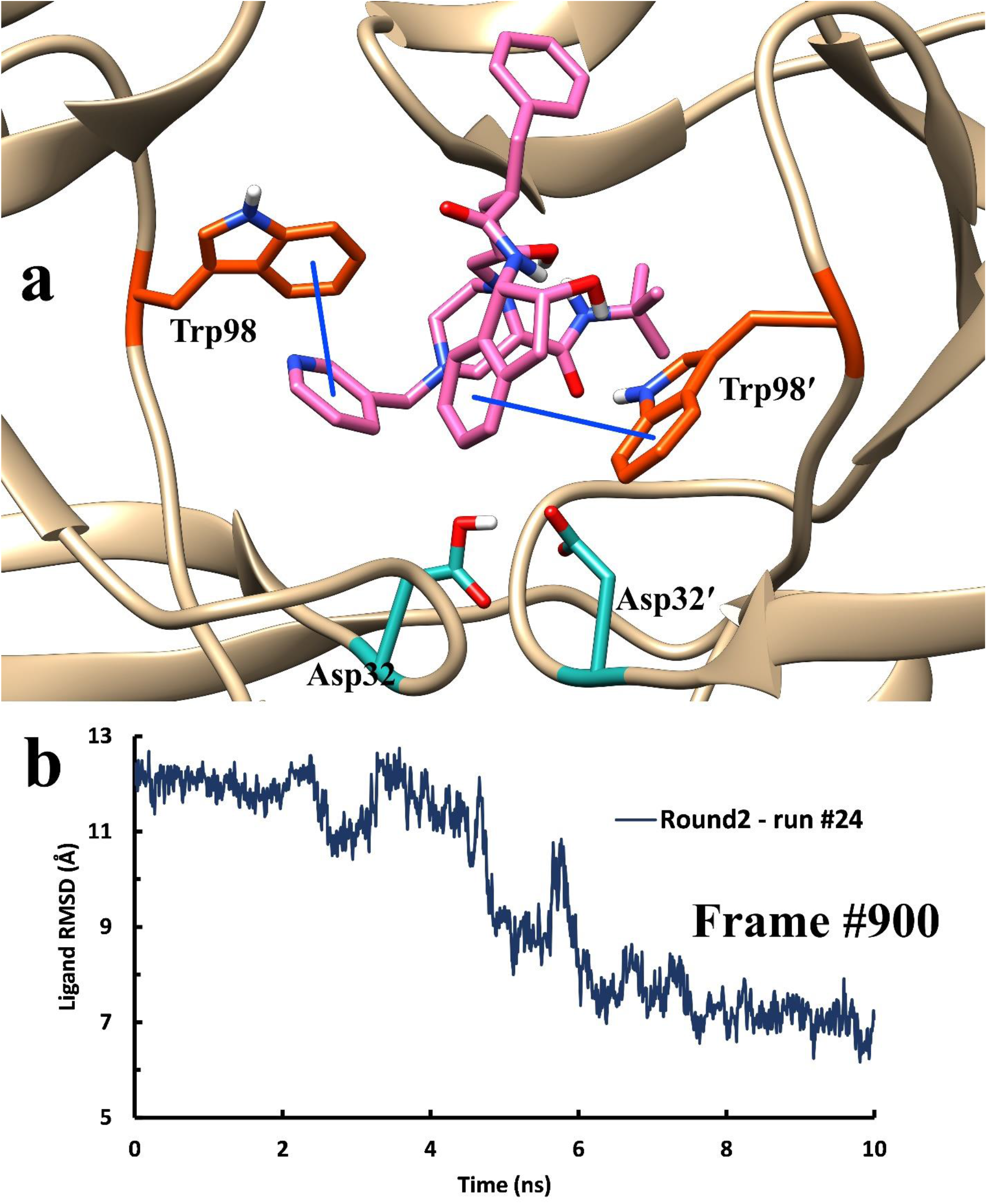
The structure of the HTLV-1 protease and the position of Indinavir at the end of the 2^nd^ stage. **a**, The position and orientation of the Indinavir molecule at the end of the entrance stage, where it interacted more with the residues in the inner parts of the binding pocket, such as Trp98 and Trp98′. The catalytic Asp residues are located in the deepest part of the binding pocket. b, The RMSD values of the Indinavir molecule in replica #24 of the second stage, in which frame #900 was chosen for the next stage.

The 3^rd^ stage was called the flattening stage #1. In this stage, significant interactions with the flap region were broken, and also the orientation of the molecule flattened, meaning that all of the chemical groups of Indinavir were roughly in the proper position (Fig. 4a). For this stage, at first, we performed 50 replicas the same as the previous stages, but no appropriate frames were found. Therefore, another 50 replicas were performed, and eventually, in replica #65, the best frame was found and selected for the next stage (Fig. 4b).

**Figure 4.**
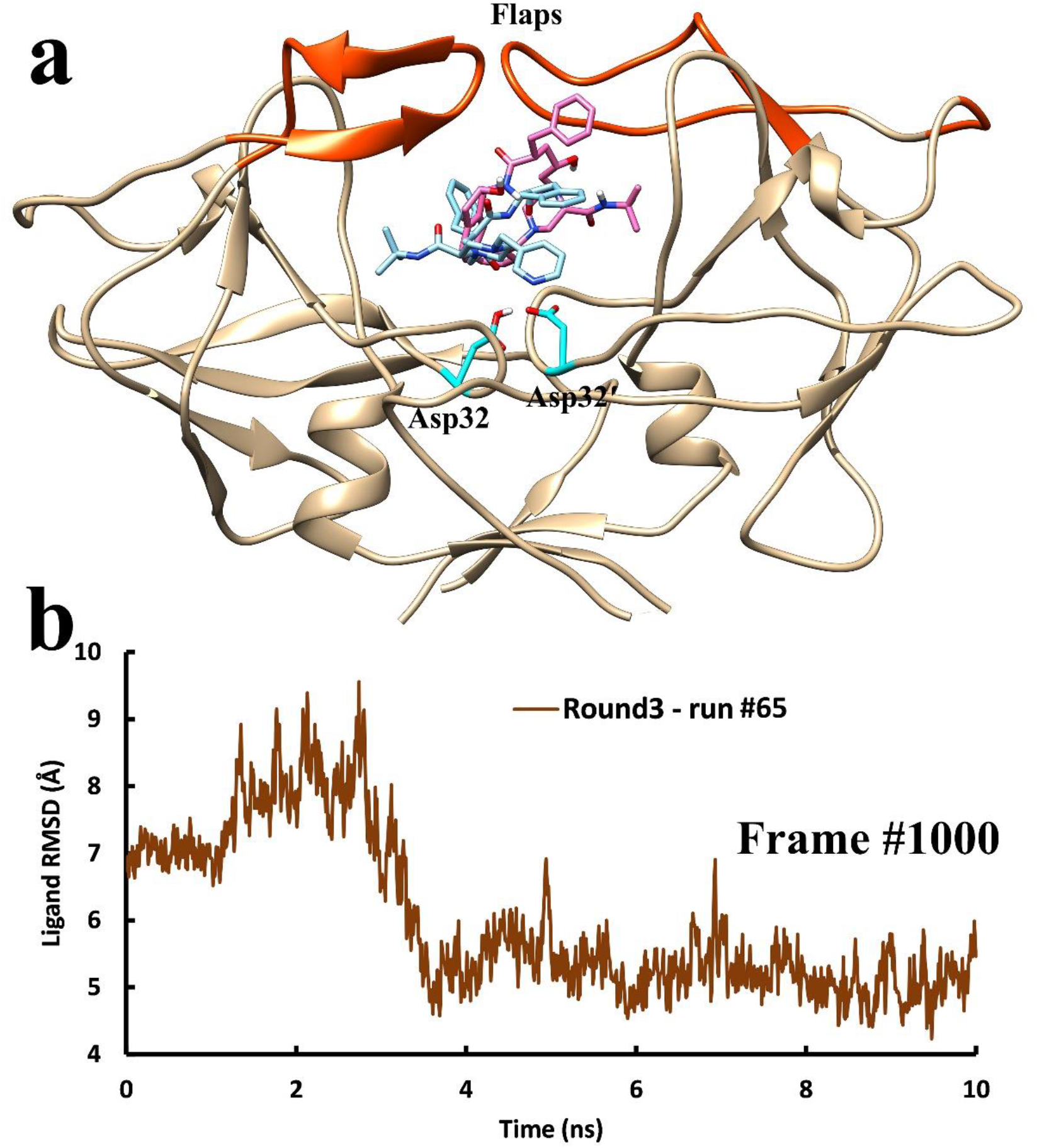
The best frames were selected from the 3^rd^ stage. **a**, The best frame was selected from replica #65 from the 3^rd^ stage, the flattening stage #1. During this stage of the binding pathway, the ligand makes fewer interactions with the flap region and interacts more with the inner parts of the binding site, such as the catalytic Asp residues. **b**, The RMSD values of Indinavir molecule in replica #65 of the 3^rd^ stage.

The 4^th^ stage was also called the flattening stage #2. It was observed that during the 10 ns of MD simulations of nearly all 100 replicas, the Indinavir molecule start to interact with the flap region and get away from the deep parts of the binding site. However, in replica #64, An appropriate conformation was found (Fig. 5a). We also monitored the ligand RMSD values of every single replica in every stage and found that in this replica, the RMSD values of the Indinavir molecule reached values below 3.5 Å (Fig. 5b). The indicated frame in replica #64 was selected as the starting point of the replicas of the next stage. In both the 3^rd^ and the 4^th^ stages, 100 replicas were performed in each stage, 200 replicas in total, to reach the desired conformation of Indinavir in the binding pocket, which indicates that this step, the flattening step, may be the rate-limiting step in the binding pathway of Indinavir to the HTLV-1 protease.

**Figure 5.**
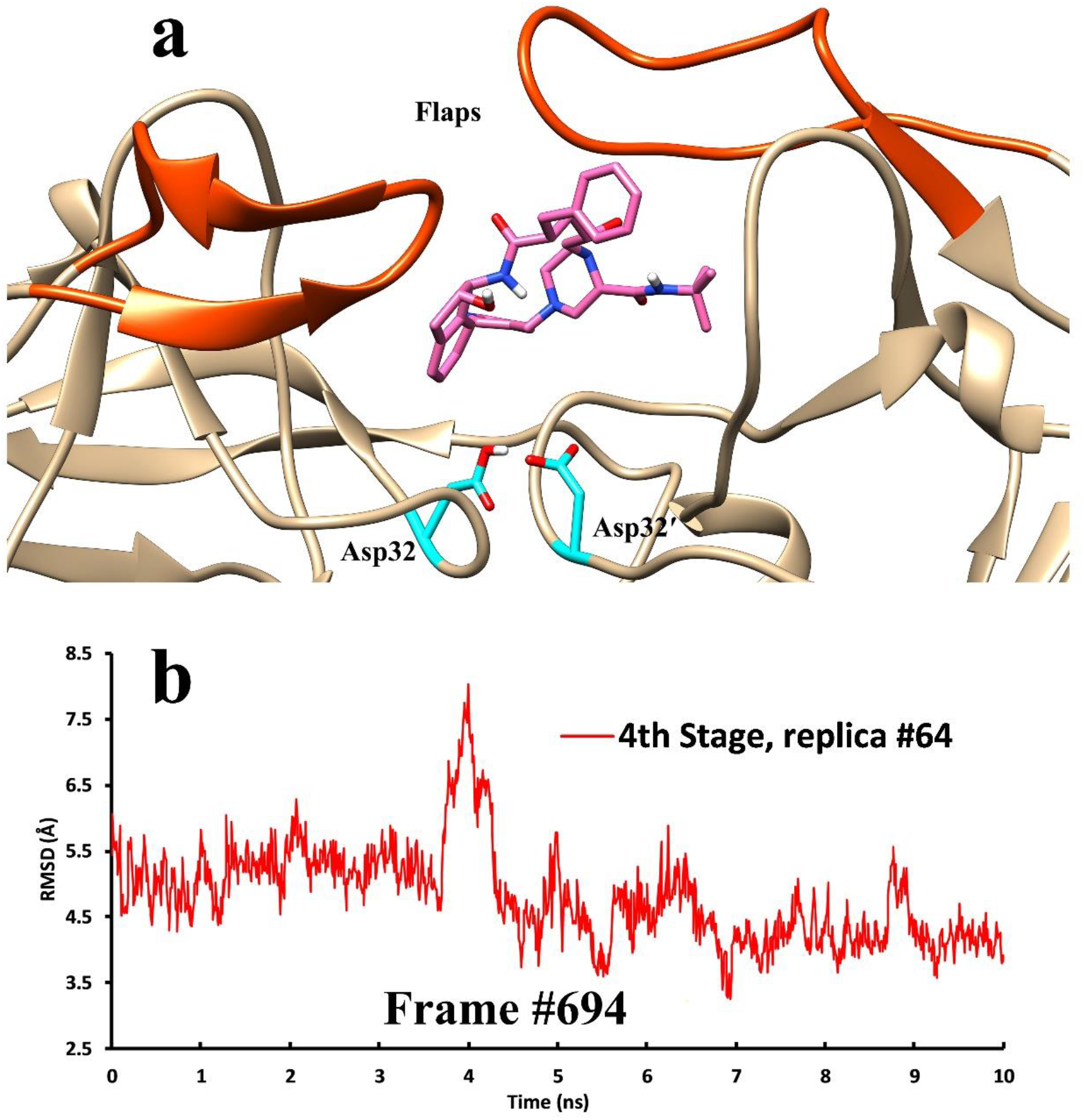
The best frames were selected from the 4th stage. a, The best frame was selected from replica #64 from the 4^th^ stage, the flattening stage #2. During this stage of the binding pathway, the ligand makes fewer interactions with the flap region and interacts more with the inner parts of the binding site, such as the catalytic Asp residues. b, The RMSD values of Indinavir molecule in replica #64 of the 4^th^ stage.

In the 5^th^ stage, the native pose stage #1, 50 replicas with a duration time of 10 ns were performed, and in replica #14, the lowest ligand RMSD values were achieved (Fig. 6b). In this stage, the significant criteria were the ligand RMSD values and forming a critical hydrogen bond between the hydroxyl group in the center of the Indinavir molecule and one of the catalytic Asp residues in the deepest part of the binding site. This information was achieved by inspecting the atomic interactions of the co-crystallized Indinavir molecule with the surrounding residues in the binding pocket of HTLV-1 protease. In our case, one of the catalytic Asp residues in the deepest part of the binding site is always protonated, which enables them to form a strong hydrogen bond with each other and avoid repulsive interactions (Fig. 6a). As mentioned, forming a hydrogen bond between Indinavir and one of these Asp residues is mandatory for Indinavir to reach the native binding pose. In replica #14, it was observed that this bond was formed, and like the previous stage, the frame with the lowest ligand RMSD value, frame #226, was chosen for the final stage, which is indicated in Fig. 6b.

**Figure 6.**
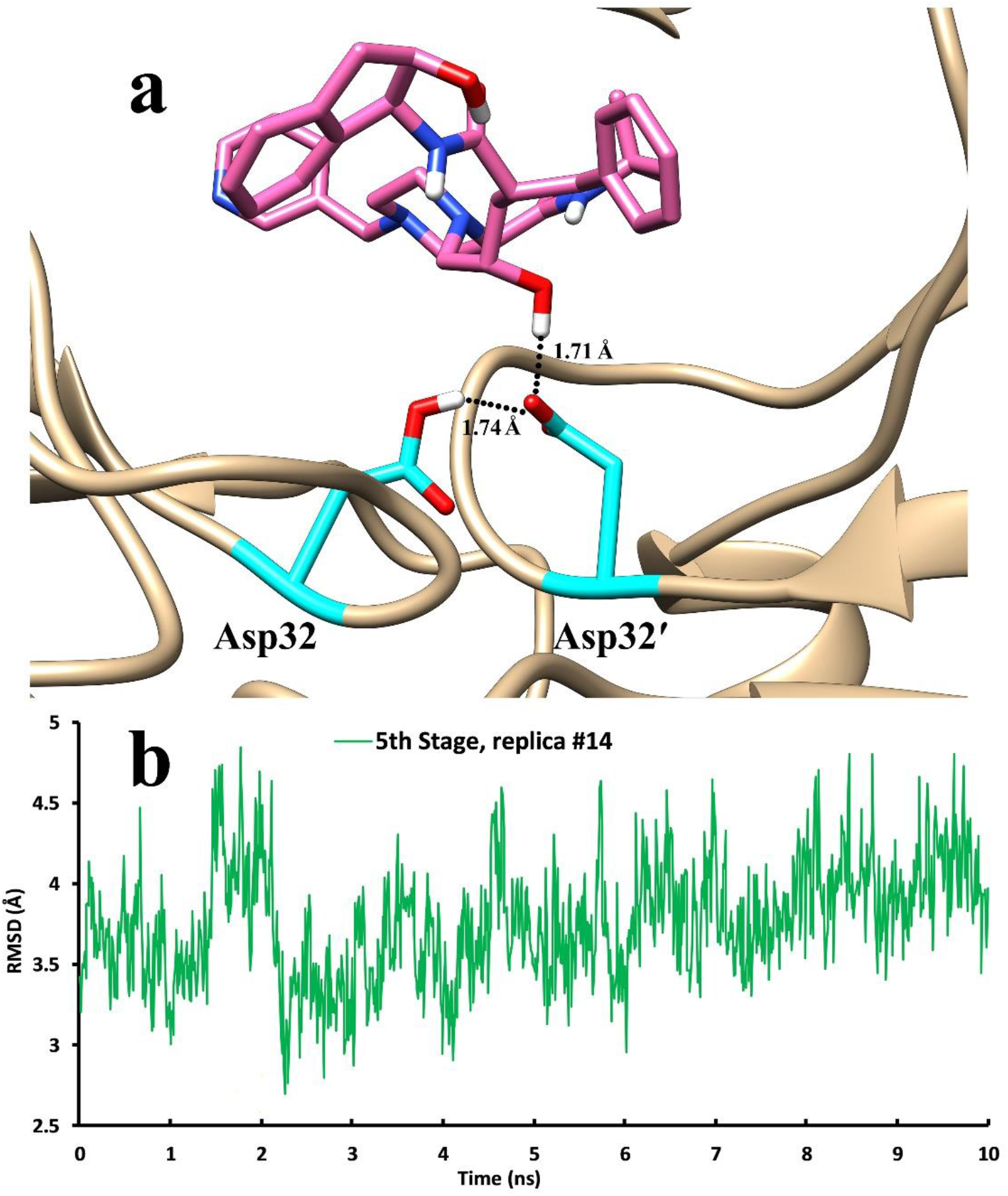
The best frame in the 5^th^ stage, the native pose stage #1. **a**, Frame #226, showing the formation of the hydrogen bond between the central Indinavir’s hydroxyl group and one of the catalytic Asp residues in the binding pocket. **b**, The RMSD values of Indinavir during the 10 ns MD simulation of replica #14 in the 5^th^ stage of the binding pathway. The best frame was chosen for the final stage is shown with an indicator.

In the final stage, the native pose stage #2, instead of performing short replicas, we performed 10 replicas with a duration time of 100 ns each. The relatively longer simulation time of the replicas in the final stage allowed the molecules to reach the most stable conformation, the native pose. Eventually, in replica #2 and #6, the native binding conformation was achieved (Fig. 7a). In this conformation, the RMSD value of the ligand was below 2.5 Å, which confirmed reaching the native binding pose (Fig. 7b). The movements and the fluctuations of the Indinavir molecule were found to be high. Despite reaching low RMSD values in the 5^th^ stage and running long MD simulations in the 6^th^ stage, it was often observed that the molecule could get out of the native binding conformation and reach higher RMSD values, as it is shown in the ligand RMSD values of the replica #6 (Fig. 7b). This may be due to the high volume of the binding site and the freedom of the Indinavir to fluctuate.

**Figure 7.**
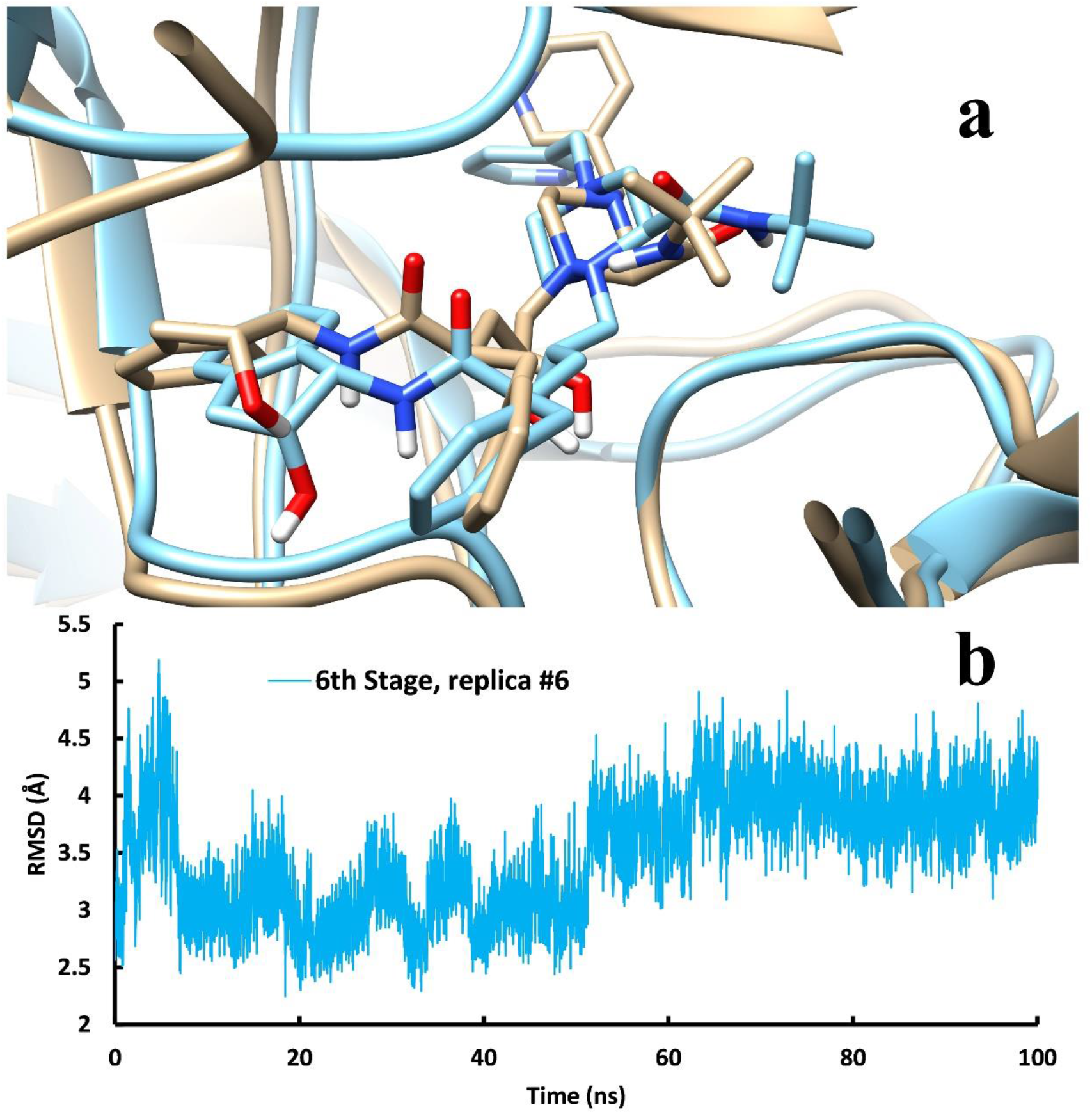
The final stage of the reconstruction of the binding pathway. **a**, The native binding conformation achieved by UUMD simulation (blue) compared to the co-crystallized structure of Indinavir inside the binding pocket of HTLV-1 protease. **b**, RMSD values of the Indinavir molecule in the 6^th^ stage replica #6 in which the ligand RMSD values reach below 2.5 Å.

Overall, in 6 stages, 360 replicas and over 4 microseconds of MD simulation were carried out to reconstruct the binding pathway of Indinavir to HTLV-1 protease (Sup Video 1). By putting successful replicas together, the binding pathway was considered as a continuous event and analyzed further. The binding pathway was about 140 ns long, in which Indinavir reached the native binding pose in just about 70 ns (Fig.8a). This showed that finding the binding site, getting inside, and reaching the native binding mode does not need extended periods and can happen in very short amounts of time. Understanding the role of different factors involved in the binding mechanism is very important. One of these crucial factors is the role of the flap region in the binding mechanism. This region has the role of first recognizing the molecule and then stabilizing it in the binding pocket. The flap region has a high fluctuation rate compared to the other region of the enzyme (Fig. 8b). This may be because this region is responsible for substrate recognition. So, it has to have high fluctuation to trap the substrate and also the required mobility to accommodate the substrate in the binding pocket. It is also responsible for stabilizing the molecule inside the pocket [35-38]. As shown in Fig. 8c, the fluctuation of the flap region is significantly lowered by the presence of the Indinavir molecule inside the binding pocket. The point indicated in the chart marks the start of the 2^nd^ stage of the MD simulation of the binding pathway. After the entrance of the molecule to the binding pocket, the flaps started to make more interactions with the indinavir molecule, which indicates the role of this region in stabilizing the molecule inside the binding pocket.

**Figure 8.**
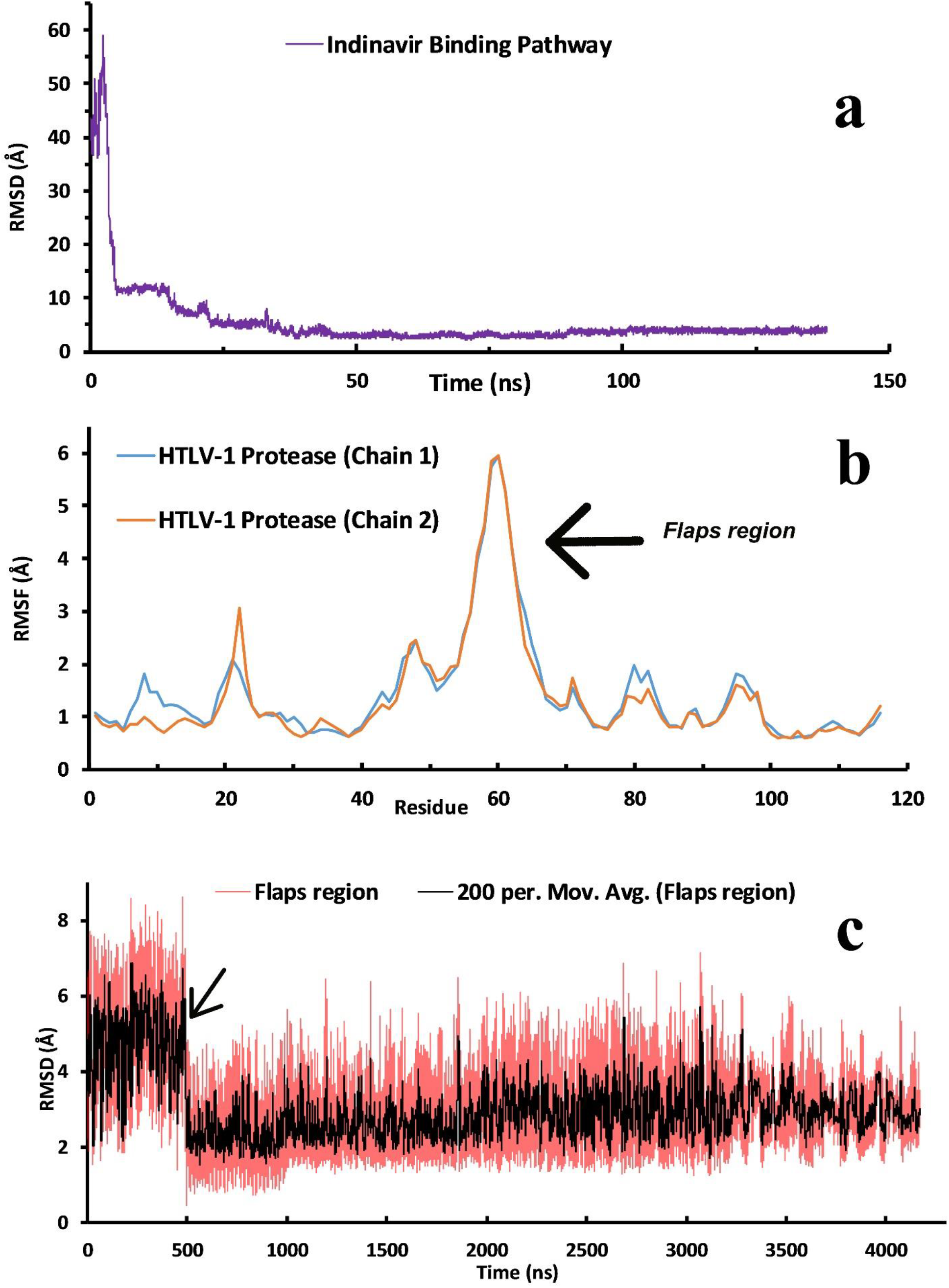
The overall analysis of the binding pathway. **a**, The RMSD values of Indinavir molecule in the binding pathway. This analysis was conducted by concatenating only all of the successful trajectories into one single trajectory file. **b**, The RMSF values of HTLV-1 protease’s residues during all of the MD simulations of the reconstruction of the binding pathway. Residues in the 50 to 70 range are considered in the flap region residues. **c**, The RMSF values of the flap region in the 4 microseconds of MD simulation performed in this study. This analysis was conducted by concatenating all of the trajectories into one single trajectory file.

During the binding pathway, the interaction energies of the molecule in the binding pathway and the contribution of each residue of the binding pocket to the interaction energies can help understand the role of essential residues and the binding pocket’s nature. Using the MMPBSA algorithm, we calculated the interaction energies, VdW and electrostatic, of the molecule during the binding pathway (Fig. 9a), and the majority of interactions are VdW interactions that reach values below -350 kJ/mol in the native binding conformation. However, the electrostatic interaction played a somewhat faded role in the binding mechanism. Another valuable analysis for understanding the role of each residue is calculating the contribution of each residue of the binding pocket to the total interaction energy (Fig. 9b). We found that the aromatic residues Trp98 and Trp98′, the catalytic residues Asp32 and Asp32′, and the flap region’s residues contribute the most to the total interaction energies. The residues from 50 to 70 are considered the residue of the flap region, which are indicated in Fig. 8b. In 2004, Kadas et al. studied the substrate specificity and sensitivity by using point mutations of the HIV and HTLV-1 proteases. They also tested the activity of Indinavir on a set of mutant forms of HTLV-1 protease and found that in some of the mutants forms, such as A59I and V56I, the Ki values of Indinavir in complex with these mutant forms were noticeably reduced [39]. This also emphasizes the importance of the residues in the flap region in the activity of the inhibitors.

**Figure 9.**
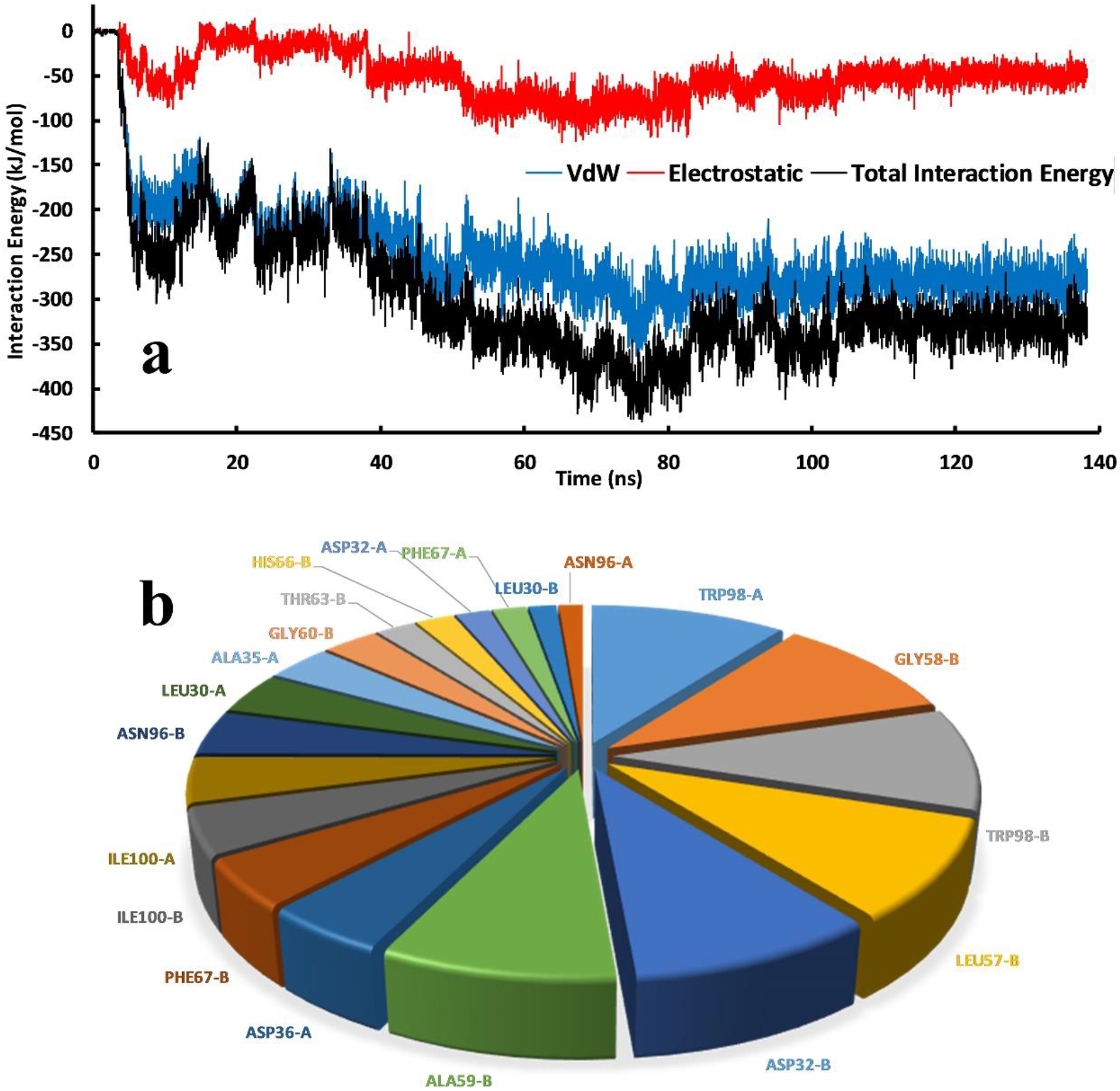
The interaction energies of the Indinavir molecule. **a**, The VdW, Electrostatic, and the total interaction energy of Indinavir molecule during the binding pathway. **b**, The contribution of the residue of the binding pocket during the binding pathway.

In the binding pathway of a ligand to its target protein, there are stable states that the ligand spends more time in them. These stable states may represent the energy barriers that the ligand has to pass in the binding pathway, and identifying them can is very important for designing more effective inhibitors. These stable states can be found in the Free energy landscape (FEL) analysis. As it is shown in Figure 10, at first, the ligand is Solvated (S) in the simulation box, and the RMSD values are high, and the contact surfaces are low, which later will be vice versa, and the RMSD values are the lowest, and the contact surfaces are the highest. This state is called the Native (N) state. Three Intermediate (I) states were found in the binding pathway between the S and N states. These intermediate states in the binding pathway may strongly be related to (i) sampling the surface of the protein and finding the binding site, (ii) engaging and interacting with the flap region, and (iii) interacting with the aromatic residues in the binding site, respectively.

**Figure 10.**
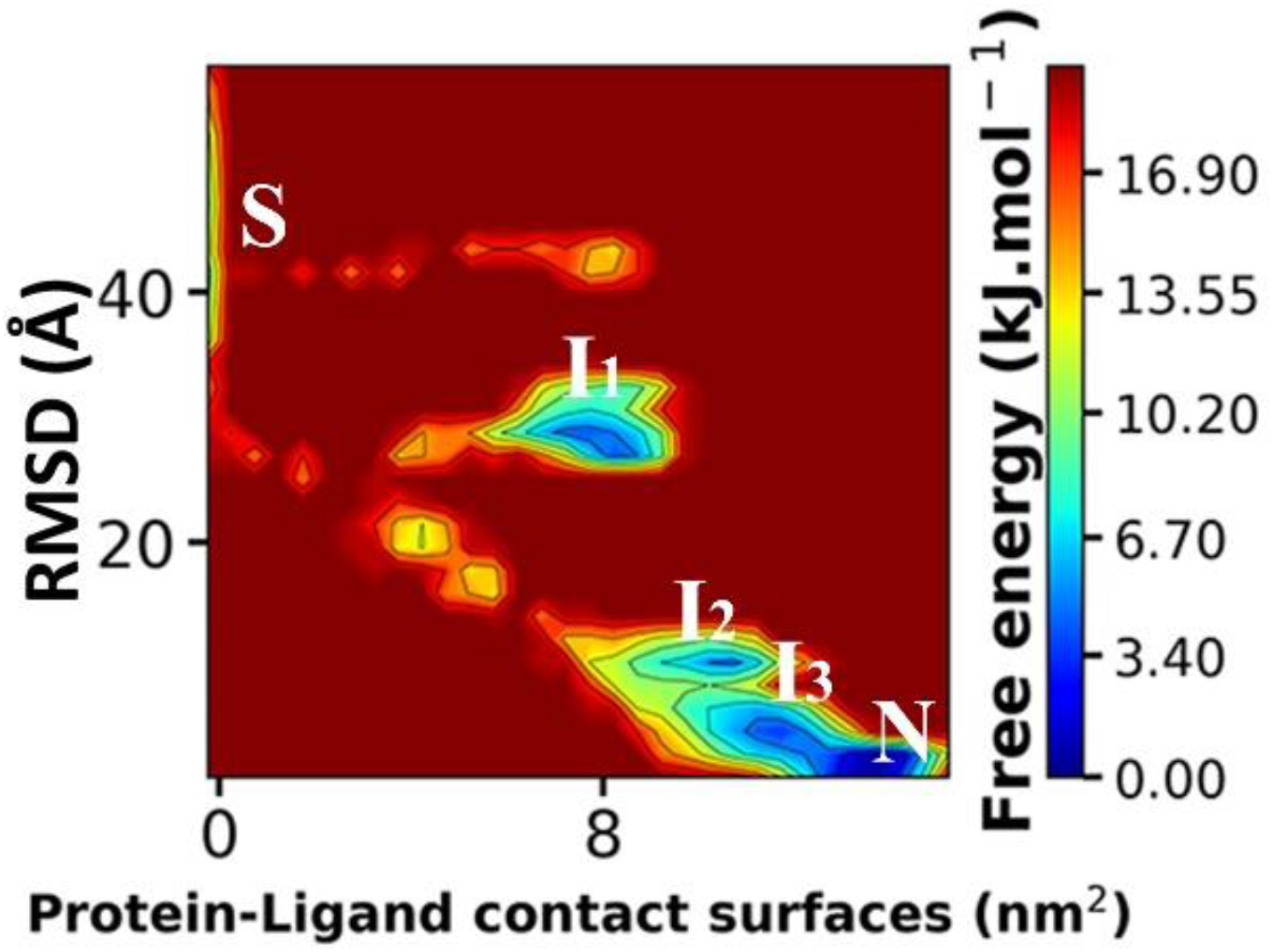
The Free Energy Landscape (FEL) of the Indinavir molecule in the binding pathway of HTLV-1 protease.

## Conclusion

In recent decades, with the outbreak of retroviruses and the death of millions of people, there is an urgent need for effective drugs to combat them. So far, effective drugs have been designed to fight viruses such as HIV, but none of these drugs has been effective against HTLV-1. One of the best molecular targets available to inhibit and control viral mechanisms is the viral protease enzyme. The use of HIV protease inhibitors has a significant healing effect on people infected with the virus. However, these HIV protease inhibitors have not shown any specific inhibitory effect on the HTLV-1 protease enzyme, and only the approved drug Indinavir has weak activity.

For this reason, we utilized UUMD simulation, which was developed by our group, to unravel the binding pathway of Indinavir to the HTLV-1 protease and its details. In this study, we characterized every step of the binding pathway by performing 360 replicas and over 4 micro-seconds of UMD simulation and found that the flap region acts as a cover for the binding pocket, and interacting and engaging with this region is a necessity for Indinavir to reach inside the binding pocket. We also found that aromatic residues such as Trp98/Trp98′ and Phe67/Phe67′, and catalytic residues Asp32/Asp32′ play an important role in the binding mechanism and have a big contribution to the interaction energies between the Indinavir molecule and the protein. In a nutshell, the details found in this study can be used for optimizing current HIV inhibitors or designing new inhibitors for the HTLV-1 protease as a strategy to fight the HTLV-1 virus, which illustrates the accuracy and the efficiency of these computational methods that can be used for such vital studies.

